# Structural and functional brain parameters related to cognitive performance across development: Replication and extension of the parieto-frontal integration theory in a single sample

**DOI:** 10.1101/659193

**Authors:** Ruben C. Gur, Ellyn R. Butler, Tyler M. Moore, Adon F.G. Rosen, Kosha Ruparel, Theodore D. Satterthwaite, David R. Roalf, Efstathios D. Gennatas, Warren B. Bilker, Russell T. Shinohara, Allison Port, Mark A. Elliott, Ragini Verma, Christos Davatzikos, Daniel H. Wolf, John A. Detre, Raquel E. Gur

**Affiliations:** Brain Behavior Laboratory and the Neurodevelopment and Psychosis Section, Department of Psychiatry, University of Pennsylvania Perelman School of Medicine, 10^th^ Floor Gates Pavilion, 3400 Spruce Street, Philadelphia, Pennsylvania 19104, USA; Department of Biostatistics, Epidemiology, and Informatics, University of Pennsylvania Perelman School of Medicine; Department of Radiology, University of Pennsylvania Perelman School of Medicine; Department of Neurology, University of Pennsylvania Perelman School of Medicine

**Author notes:** Corresponding author Ruben C. Gur, PhD, Brain Behavior Laboratory, 10^th^ Floor Gates Pavilion/HUP, 3400 Spruce Street, Philadelphia, PA 19104.

**Keywords:** Neuroimaging, multimodal brain parameters, brain performance relation, neurodevelopment, neurocognition

## Abstract

The Parieto-Frontal Integration Theory (PFIT) identified a fronto-parietal network of regions where individual differences in brain parameters most strongly relate to cognitive performance. PFIT was supported and extended in adult samples, but not in youths or within single-scanner well-powered multimodal studies. We performed multimodal neuroimaging in 1601 youths age 8-22 on the same 3-Tesla scanner with contemporaneous neurocognitive assessment, measuring volume, gray matter density (GMD), mean diffusivity (MD), cerebral blood flow (CBF), resting-state functional MRI measures of amplitude of low frequency fluctuations (ALFF) and regional homogeneity (ReHo), and activation to a working memory and a social cognition task. Across age and sex groups, better performance was associated with higher volumes, greater GMD, lower MD, lower CBF, higher ALFF and ReHo and greater activation for the working memory task in PFIT regions. However, additional cortical, striatal, limbic and cerebellar regions showed comparable effects, hence PFIT needs expansion into an Extended PFIT (ExtPFIT) network incorporating nodes that support motivation and affect. Associations of brain parameters became stronger with advancing age group from childhood to adolescence to young adulthood, effects occurring earlier in females. This ExtPFIT network is developmentally fine-tuned, optimizing abundance and integrity of neural tissue while maintaining low resting energy state.

---

While multiple parameters of brain structure and function have been examined with structural and functional magnetic resonance imaging studies, it is still unclear how these measures are related to arguably the main product of brain processes - cognitive performance. The relation between brain volume and cognitive performance has received the most extensive investigation since Frederick Tiedemann (1836) offered his conclusion that “There is undoubtedly a very close connexion between the absolute size of the brain and the intellectual powers and functions of the mind.” (p. 502). However, the magnitude of this relation is still debated; estimates of variance in cognitive measures explained by volume range from -3% to >30% (Witelson, Beresh, & Kigar, 2005; Gignac and Bates, 2017; Nave et al. 2018; Pietschnig et al. 2015). Advanced neuroimaging offers additional parameters of brain structure and function, and Haier and Jung (2007) performed a review of studies across modalities and proposed the Parieto-Frontal Integration Theory (PFIT), which stipulates a network of regions that are predominantly involved in complex reasoning and intelligence tasks. This network integrates dorsolateral prefrontal cortex, the inferior and superior parietal lobule, the anterior cingulate, and regions within the temporal and occipital lobes (Basten, Hilger and Fiebach 2015). Within this network of regions, neuroanatomic parameters of higher volume, density and anisotropy and lower rates of cerebral blood flow and metabolism have been associated with better cognitive performance.

The PFIT theory received some support in subsequent studies, although few have examined multiple brain parameters typically focusing only on volume. Ritchie et al. (2015)compared the relationship of volume to performance with other neuroanatomic parameters such as cortical thickness and found that volume accounted for the largest share of the variance (around 12%). Ryman et al. (2016) applied graph-theory analyses to volumetric data and reported that in males a latent factor of fronto-parietal gray matter related to general cognitive abilities, while in females the cognition-related factor involved white matter efficiency and total gray matter volume without regional specificity. Studies with relatively small samples have linked some diffusion tensor imaging (DTI) parameters to performance (e.g., Schmithorst, Wilke, Dardzinski, Holland 2005; Qiu, Tan, Zhou and Khong 2008; Gen^ et al. 2018). A study on a larger sample (N=72) applied graph theory to DTI data and reported sex differences, with females having greater local efficiency, but these effects were not linked to performance (Yan et al. 2011). Graph theory was also applied in the PNC sample by Ingalhalikar et al (2014), and they reported greater within-hemispheric connectivity in males and between-hemispheric connectivity in females, as well as sex differences in modularity and participation coefficients.

Studies relating cerebral blood flow (CBF) to performance likewise supported the PFIT model, showing that age-related decline in CBF is related to performance decline (Hshieh et al. 2017, Rane et al. 2018). Resting-state functional MRI (rs fMRI) was examined in relation to cognitive performance in a small sample (Pamplona et al. 2015), and in a larger sample (N=79) by Vakhtin et al. (2014), who assessed both resting state and task-activated connectivity. They reported that regions involved in task related networks included bilateral medial frontal and parietal cortex, right superior frontal lobule, and right cingulate gyrus. As part of multivariate measures in the Human Connectome Project (Smith et al. 2015), Finn et al. (2015) showed that functional connectivity profiles predicted “levels of fluid intelligence” and that “the same networks that were most discriminating of individuals were also most predictive of cognitive behavior.” Furthermore, Yoo et al. (2018) reported, across several data sets, that models trained on task data outperformed those trained on resting state data in predicting performance (cf. Fong et al. 2019, Jangraw et al. 2018) and Greene, Gao, Scheinost, & Constable (2018) demonstrated similar effects in two large data sets. Finally, Dubois, Galdi, Paul, & Adolphs (2018) were able to predict up to 20% of variance in general cognitive performance based on resting-state connectivity metrics in a large sample (n=884) from the HCP. Each of these studies examined individual parameters of either structure or function. Therefore, there is need for simultaneous examination of neuroanatomic and neurophysiologic parameters in relation to cognitive performance. Such simultaneous examination will allow gauging the relative contribution of the anatomic and physiologic parameters to cognitive performance, and their examination during development will offer insight on how brain parameters are fine-tuned for optimal adult levels. The PFIT has yet to be tested across both structural and functional brain parameters in a single, adequately powered multimodal study of youths.

We tested the PFIT with multimodal neuroimaging in a prospective sample of 1601 youths age 8 to 22, all studied on the same high-field (3 Tesla) scanner with contemporaneously obtained measures of cognitive performance, as part of the Philadelphia Neurodevelopmental Cohort (Gur et al. 2012; Calkins et al. 2015). The methods of sample ascertainment and the detailed neuroimaging protocols have been published (Satterthwaite et al. 2014a). Multimodal neuroimaging yielded regional measures of gray matter (GM) and white matter (WM) volume and GM density (GMD) from T1-weighted scans, mean diffusivity (MD) from diffusion tensor imaging (DTI; fractional anisotropy was also measured but since it is only valid in white matter, this parameter was not included here), resting state cerebral blood flow (CBF) from arterial spin-labeled (ASL) sequences, amplitude of low frequency fluctuations (ALFF) and regional homogeneity (ReHo) measures from rs fMRI, and BOLD activation for a working-memory (NBack) and a social-cognition (Emotion identification; IDEmo) task. The methods for image processing and for obtaining these brain parameters were detailed in previous publications (Gennatas et al. 2017, Ingalhalikar et al. 2014, Satterthwaite et al. 2014a,b), and are briefly summarized in Methods below. The neurocognitive assessment provided measures of accuracy and speed on multiple behavioral domains. Since available literature primarily examined general intellectual functioning, we selected as the primary cognitive measure the most comparable score from the battery, which is a factor score that summarizes accuracy on executive functioning and complex cognition (Moore et al. 2015, Swagerman et al. 2016).

## MATERIALS AND METHODS

### Participants

Participants for the PNC were recruited from the Children’s Hospital of Philadelphia (CHOP) pediatric network throughout the Delaware Valley as described in Calkins et al. (2015). A subsample of 1,601 participants (out of the 9498 PNC sample) underwent multimodal neuroimaging as described in Satterthwaite et al. (2014a). Of these, 340 were excluded for medical disorders that could affect brain function, as well as current use of psychoactive medications, prior inpatient psychiatric treatment, or an incidentally encountered structural brain abnormality. Sample size was further reduced for some modalities upon quality assurance procedures, most for excessive motion (see below, Table 1 and Supplementary Table SI). All participants underwent psychiatric assessment (Calkins et al. 2015) and neurocognitive testing (Guret al. 2012, 2015).

**Table 1.**
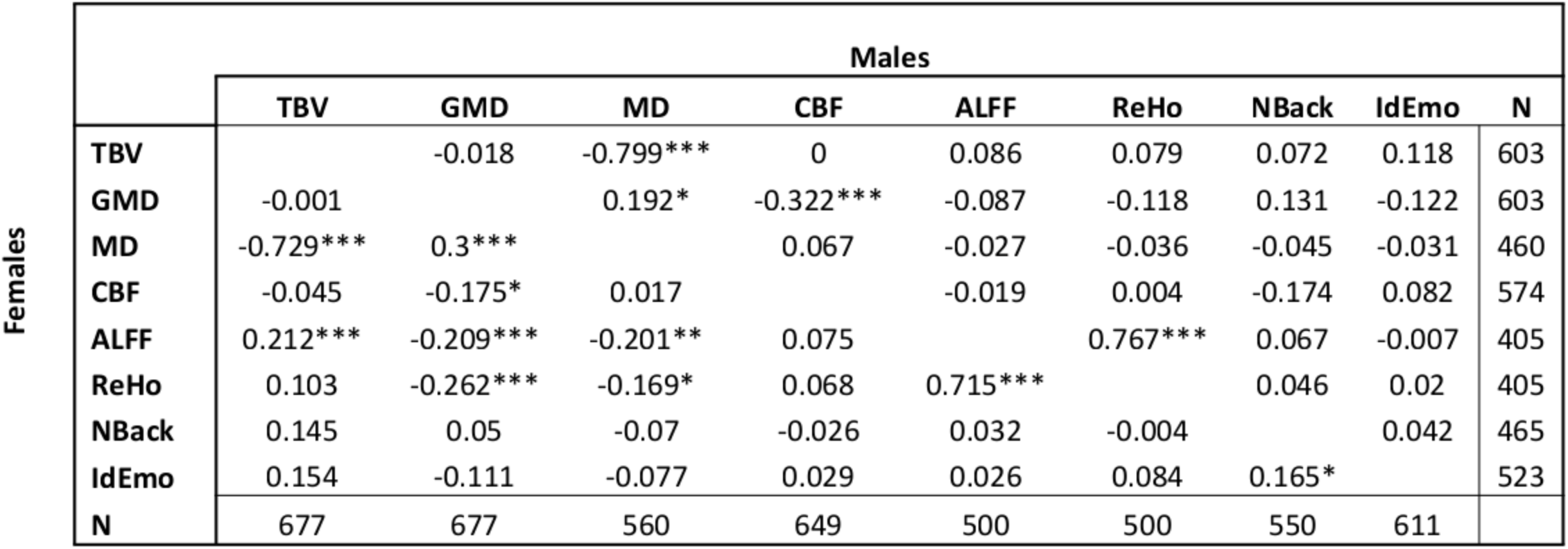
Intercorrelations among the global brain parameters for females (upper triangle) and males (lower triangle). * p < .05; ** p < .01, *** p < .001. TBV=Total brain volume; GMD=Gray matter density; MD=Mean diffusivity; CBF=Cerebral blood flow; ALFF= Amplitude of low frequency fluctuations at resting-state; ReHo= Regional homogeneity at resting-state; Nback=Blood oxygenation-level dependent (BOLD) activation for the Nback task; IDEmo=BOLD activation for the emotion identification task.

### Cognitive measures

Cognitive performance was assessed with the Penn Computerized Neurocognitive Battery (CNB). The CNB consists of 14 tests adapted from tasks applied in functional neuroimaging to evaluate a range of cognitive domains (Gur et al. 2010, 2012, 2014; Moore et al. 2015; Roalf et al. 2014). These domains include executive control (abstraction and mental flexibility, attention, working memory), episodic memory (verbal, facial, spatial), complex cognition (verbal reasoning, nonverbal reasoning, spatial processing), social cognition (emotion identification, emotion intensity differentiation, age differentiation) and sensorimotor and motor speed. Accuracy and speed for each test were z-transformed. Cognitive performance was summarized by a factor analysis of both speed and accuracy data (Moore et al. 2015), which delineated three accuracy factors corresponding to: 1) executive function and complex reasoning, 2) episodic memory, and 3) social cognition. The first factor was used as the measure of cognitive performance in all analyses, as it has the highest association with IQ estimates (Moore et al 2015; Swagerman et al. 2016).

### Neuroimaging

All MRI scans were acquired on a single 3T Siemens TIM Trio whole body scanner located in the Hospital of the University of Pennsylvania. Signal excitation and reception were obtained using a quadrature body coil for transmit and a 32-channel head coil for receive. Gradient performance was 45 mT/m, with a maximum slew rate of 200 T/m/s. Image processing and analysis was performed using AFNI, FSL, and DTI specific tools with the Advanced Neuroimaging Tools (ANTs) pipeline (Avants et al. 201 la,b).

#### Structural

Parameters of brain anatomy were derived from volumetric scans (T1-weighted) and from diffusion tensor imaging.

##### Volumetric MRI

Brain volumetric imaging was obtained using a magnetization prepared, rapid-acquisition gradient-echo (MPRAGE) sequence (TR/TE/TI=1810/3.5/1100ms; FOV RL/AP=180/240mm; Matrix RL/AP/slices=192/256/160 Slice thick/gap=l/0mm; Flip angle=9°; No Reps; GRAPPA factor=2; BW/pixel=130 Hz; PE direction=RL; Acq time=3:28 min). Receive coil shading was reduced by selecting the Siemens prescan normalize option, which is based on a body coil reference scan. Image quality assessment (QA) was performed both by visual inspection and with algorithms to detect artifacts such as related to excessive head motion.

To maximize accuracy, advanced structural image processing, quality assurance and registration procedures were employed for measurement of the cortical subcortical and cerebellar volumes and gray matter density. Estimation of brain regions used a multi-atlas labeling approach. A set of 24 young adult T1-weighted volumes from the OASIS data set (Marcus et al. 2007) were manually labeled and registered to each subject’s T1-weighted volume using the top-performing SyN diffeomorphic registration (Avants et al. 201 la; Klein et al. 2010). These label sets were synthesized into a final parcellation using joint label fusion, which is similarly reliable to other state-of-the art label fusion algorithms but uses significantly fewer atlases, and is far more accurate than segmentation performed with a single atlas (Wang et al. 2013). Volume was determined for each parcel using the intersection between the parcel created and prior driven gray matter cortical segmentation from the ANTs cortical thickness pipeline as described below. Density estimates were calculated within each parcel as described below. To avoid registration bias and maximize sensitivity to detect regional effects that can be impacted by registration error, a custom adolescent template and tissue priors were created using data from 140 PNC participants, balanced for age and sex. Structural images were then processed and registered to this custom template using the ANTs cortical thickness pipeline (Tustison et al. 2014). This procedure includes brain extraction, N4 bias field correction (Tustison et al. 2010), Atropos tissue segmentation (Avants et al. 201 lb), and SyN diffeomorphic registration method (Avants et al. 2011a; Klein et al.2010).

Finally, gray matter density was calculated using Atropos (Avants et al. 201 lb), with an iterative segmentation procedure that is initialized using 3-class K-means segmentation. This procedure produces both a discrete 3-class hard segmentation as well as a probabilistic gray matter density map (soft segmentation) for each subject. Gray matter density (GMD) was calculated within the intersection of this 3-class segmentation and the subject’s volumetric parcellation (Gennatas et al. 2017). Images included in the final analysis passed a rigorous QA procedure, including evaluation of motion, as previously detailed (Rosen et al. 2018).

##### Diffusion (DTI)

Diffusion weighted imaging (DWI) scans for measuring water diffusion were obtained using a twice-refocused spin echo (TRSE) single-shot EPI sequence. The sequence employs a four lobed diffusion encoding gradient scheme combined with a 90-180-180 spin-echo sequence designed to minimize eddy-current artifacts. The sequence consisted of 64 diffusion-weighted directions with b = 1000 s/mm^2^, and 7 scans with b = 0 s/mm^2^.

Diffusion data were skull stripped by generating a brain mask for each subject by registering a binary mask of a standard image (FMRIB58 FA) to each subject’s brain using FLIRT (Jenkinson et al. 2002). When necessary, manual adjustments were made to this mask. Next, eddy currents and movement were estimated and corrected using FSL’s eddy tool (Andersson and Sotiropoulos, 2016; Graham et al. 2016; Roalf et al. 2016). Eddy improves upon FSL’s Diffusion Tool Box (Behrens et al. 2003) and eddy correct tool (Andersson and Sotiropoulos 2016; Graham et al. 2016) by simultaneously modeling the effects of diffusion eddy current and head movement on DTI images, reducing the amount of resampling. The diffusion gradient vectors were rotated to adjust for motion using the 6-parameter motion output generated from eddy. Then, the BO field map was estimated and distortion correction was applied to the DTI data using FSL’s FUGUE (Smith 2002). Finally, the diffusion tensor was modeled and metrics (MD) were estimated at each voxel using FSL’s DTIFIT.

Registration from native space to a template space was completed using DTI-TK (Zhang et al. 2014; Zhang et al. 2006). First, DTI output files from DTIFIT were converted to DTI-TK format. Next, a template was generated from the tensor volumes using 14 representative diffusion data sets that were considered “Excellent” from the PNC sample. One individual from each of the 14 ages (age range 8-21) was randomly selected. These 14 DTI volumes were averaged to create an initial template. Next, data from the 14 subjects were registered to this template in an iterative manner. Unlike standard intensity-based registration algorithms, this process utilizes the full tensor information to best align the underlying white matter tracts using iterations of rigid, affine and diffeomorphic registration leading to the generation of a successively refined template. Ultimately, one high-resolution refined template was created and used for registration of the remaining diffusion datasets. All DTI maps were then registered (rigid, affine, diffeomorphic) to the high-resolution study-specific template using DTI-TK.

Whole brain analysis was performed using a customized implementation of tract-based spatial statistics (TBSS) (Bach et al. 2014). MD values were computed using a study specific white matter skeleton. MD has excellent intrasession and acceptable intersession reliability, especially for sequences with more gradient directions and repetitions within a session, as is the case in the present study (Wang et al. 2012). Then, standard regions of interest (ROI; ICBM-JHU Whit Matter Tracts; Harvard-Oxford Atlas) were registered from MNI152 space to the study-specific template using ANTs registration (Avants et al. 2011). Mean diffusion metrics were extracted from these ROIs using FSL’s ‘fslmeants’. Images included in this final analysis passed a stringent quality assessment procedure as previously detailed (Roalf et al. 2016).

#### Functional

##### Perfusion (ASL) measures of cerebral blood flow (CBF)

Brain perfusion was imaged using a pseudo continuous arterial spin labeling (pCASL) sequence (Wu et al. 2007), which has been shown to have good to excellent scan-rescan reliability within site (Almeida et al. 2018). The sequence used a single-shot spin-echo EPI readout. Parallel acceleration (i.e. GRAPPA factor = 2) was used to reduce the minimum achievable echo time. The arterial spin labeling parameters were: label duration = 1500ms, post-label delay = 1200ms, labeling plane = 90mm inferior to the center slice. The sequence alternated between label and control acquisitions for a total of 80 acquired volumes (40 labels and 40 controls), the first being a label.

ASL data were pre-processed using standard tools included with FSL (Jenkinson et al. 2012). Following distortion correction using the B0 map with FUGUE, the first four image pairs were removed, the time series was realigned in MCFLIRT (Jenkinson et al. 2002), the skull was removed with BET (Smith 2002), and the image was smoothed at 6mm FWHM using SUSAN (Smith and Brady 1997). CBF was quantified from control-label pairs using ASL Toolbox (Wang et al. 2008). As prior (Satterthwaite et al. 2014a), the T1 relaxation parameter was modeled on an age- and sex-specific basis (Wu et al. 2010). This model accounts for the fact that T1 relaxation time differs according to age and sex, and has been shown to enhance the accuracy and reliability of results in developmental samples (Jain et al. 2012). The CBF image was coregistered to the T1 image using boundary-based registration (Greve and Fischl 2009), and regional CBF values were averaged within each parcel. Subjects included in this analysis had low motion as measured by mean relative framewise displacement less than 0.5mm.

##### Resting-state BOLD

Resting-state BOLD scans were acquired with a single-shot, interleaved multi-slice, gradient-echo, echo planar imaging (GE-EPI) sequence. In order to reach steady-state signal levels, the sequence performed two additional dummy scans at the start. The imaging volume was sufficient to cover the entire cerebrum of all subjects, starting superiorly at the apex. In some subjects, the inferior portion of the cerebellum could not be completely included within the imaging volume. The selection of imaging parameters was driven by the goal of achieving whole brain coverage with acceptable image repetition time (i.e. TR = 3000ms). A voxel resolution of 3 * 3* 3mm with 46 slices was the highest obtainable resolution that satisfied those constraints (multiband acquisition methods were not available when this data was collected). During the resting-state scan, a fixation cross was displayed as images were acquired. Participants were instructed to stay awake, keep their eyes open, fixate on the displayed crosshair, and remain still. Resting state scan duration was 6.2min.

##### Task-related BOLD

Task-related modulations of BOLD were measured using two fMRI tasks as described in Satterthwaite et al. 2014. The Nback working memory (WM) task involved presentation of complex geometric figures (fractals) for 500ms, followed by a fixed interstimulus interval of 2500ms. This occurred under three conditions: 0-back, l-back, and 2-back, producing different levels of WM load. In the 0-back condition, participants responded with a button press to a specified target fractal. For the 1-back condition, participants responded if the current fractal was identical to the previous one; in the 2-back condition, participants responded if the current fractal was identical to the item presented two trials previously. Each condition consisted of a 20-trial block (60s); each level was repeated over three blocks. The target-foil ratio was 1:3 in all blocks, with 45 targets and 135 foils overall. Visual instructions (9s) preceded each block, informing the participant of the upcoming condition. The task included a total of 72s of rest while a fixation crosshair was displayed, which was distributed equally in three blocks of 24s at beginning, middle, and end of the task. Total working memory task duration was 11,6min. The emotion identification (IdEmo) task employed a fast event-related design with a jittered inter-stimulus interval (ISI). Participants viewed 60 faces displaying neutral, happy, sad, angry, or fearful expressions, and were asked to label the emotion displayed. Each face was displayed for 5.5s followed by a variable ISI of 0.5 to 18.5s, during which a complex crosshair (that matched the faces’ perceptual qualities) was displayed. Total IdEmo task duration was 10.5min.

##### BOLD Processing

Both task-related and resting-state task-free functional images were processed using a top-performing pipeline for removal of motion-related artifact (Ciric et al. 2017). Preprocessing steps included (1) correction for distortions induced by magnetic field inhomogeneities using FSL’s FUGUE utility, (2) removal of the 4 initial volumes of each acquisition, (3) realignment of all volumes to a selected reference volume using MCFLIRT (Jenkinson et al. 2002), (4) removal of and interpolation over intensity outliers in each voxel’s time series using AFNI’s 3DDESPIKE utility (Cox 1996), (5) demeaning and removal of any linear or quadratic trends, and (6) co-registration of functional data to the high-resolution structural image using boundary-based registration (Greve and Fischl 2009). The artifactual variance in the data was modelled using a total of 36 parameters, including the 6 framewise estimates of motion, the mean signal extracted from eroded white matter and cerebrospinal fluid compartments, the mean signal extracted from the entire brain, the derivatives of each of these 9 parameters, and quadratic terms of each of the 9 parameters and their derivatives. Both the BOLD-weighted time series and the artifactual model time series were temporally filtered using a first-order Butterworth filter with a passband between 0.01 and 0.08 Hz.

Subject exclusions were based on BOLD sequence specific quality control assessment. These included: lack of complete fMRI data for a particular task; high in-scanner motion (mean relative displacement >0.5 mm or maximum relative displacement >6mm); poor brain coverage in fMRI data; or failure to perform tasks at a minimal level (N-back: More than 8 (>2 SD) non-responses on the 0-back; Emotion Identification: More than 11 (>2 SD) non-responses, or number correct not significantly above chance performance [<18]).

##### BOLD Outcome measures

###### ALFF

Functional connectivity among brain regions is primarily attributable to correlations among low-frequency fluctuations in regional activation patterns (Di et al. 2013). The amplitude of low frequency fluctuations shows high scan-rescan reliability within a session, with ICCs centered at approximately .8 (Somandepalli et al. 2015). The voxelwise amplitude of low-frequency fluctuations (ALFF; Zang et al. 2007) was computed as the sum (discretised integral) over frequency bins in the low-frequency (0.01-0.08Hz) band of the voxelwise power spectrum, computed using a Fourier transform of the time-domain of the voxelwise signal. ALFF was calculated on data smoothed in SUSAN using a Gaussian-weighted kernel with 6mm FWHM (Smith and Brady 1997). ALFF was selected because it is a frequently used measure derived from fMRI (PubMed crossing of fMRI and ALFF yielded 597 publications https://www-ncbi-nlm-nih-gov.proxy.library.upenn.edu/pubmed/?term=ALFF+and+fmriretrieved3/8/2020).

###### ReHo

Voxelwise regional homogeneity (ReHo; Zang et al. 2004) is equivalent to Kendall’s coefficient of concordance computed over the timeseries in each voxel’s local neighborhood. ReHo can thus be used as an estimate of the homogeneity of each neighborhood’s activation pattern. ReHo shows moderate scan-rescan reliability within a session, with ICCs centered at approximately .6 (Somandepalli et al. 2015). Because spatial smoothing intrinsically elevates ReHo estimates by elevating spatial autocorrelation, Kendall’s W was computed only on unsmoothed data. Each voxel’s neighborhood was defined to include the 26 voxels adjoining its faces, edges, and vertices. The voxelwise homogeneity map was subsequently smoothed using a Gaussian kernel with FWHM of 6mm in SUSAN to improve the signal-to-noise ratio (Smith and Brady 1997). Finally, regional ReHo values were then averaged across the anatomically derived subject specific segmentation. Participants included in this analysis had low motion with mean relative frame wise displacement less than 2.5mm. ReHo was selected because it is among the most frequently used measures derived from resting-state fMRI (PubMed crossing of fMRI and ReHo yielded 549 manuscripts https://www-ncbi-nlm-nih-gov.proxy.library.upenn.edu/pubmed/?term=reho+and+fmriretrieved3/8/2020).

ALFF and ReHo reflect different aspects of regional neural activity. ALFF measures the total power of the BOLD signal within the low-frequency range, and is thus proportional to regional neural activity, while ReHo is a voxel-based measure of the similarity between the timeseries of a given voxel and its nearest neighbors, reflecting the synchrony of adjacent regions (see Lv et al. 2018, Zang et al. 2004).

Subject-level statistical analyses were carried out voxelwise with a canonical hemodynamic response function in FSL FEAT. For the n-back, three condition blocks (0-back, 1-back, and 2-back) were modeled. Six motion parameters and the instruction period were included as nuisance covariates, and the rest (fixation) condition provided unmodeled baseline. The dependent measure obtained was 2-back > 0-back, capturing the effect of increasing working memory load. For emotion identification, events were modeled as 5.5sec boxcar, matching the duration of face presentation. Five individual emotion regressors were included together with their temporal derivatives and six motion parameters. The contrast of interest was Emotion Face (happy+sad+anger + fear+neutral> fixation). Target ROIs for an NBack and an emotion identification task have acceptable reliability, as measured by relative agreement ICCs, over the course of two weeks (Plitchta et al. 2012; but see Elliott et al. 2020).

### Statistical analysis

We examined the association of variability in brain structural and functional parameters and our performance parameter in four stages, aimed to contain type I error: 1. Global values were examined for association with performance; 2. Hypothesis testing, specifically the PFIT regions were examined across modalities compared to non-PFIT counterparts; 3. Exploratory (hypothesis-generation), all regions were included and non-PFIT regions showing comparable significance and effect sizes were considered for inclusion in an Extended-PFIT (ExtPFIT) network; 4. Data-driven performance prediction to estimate variance explained by brain parameters. Prior to further analyses, we compared correlations between performance and brain parameters in the left and right hemispheres, and found that the effect sizes separating high and low performers were nearly identical in the two hemisphere for given regions, with left-right correlations ranging from the lowest of .833 for CBF to .944 for ReHo. Therefore, subsequent analyses summed or volume-averaged the two hemispheres as appropriate. Analyses were conducted using the open source R platform (Version 3.5, R Core Team 2015).

#### 1. Global values

The global measures included estimated total brain volume, average whole-brain values of GMD, MD, CBF, ALFF, ReHo, and the BOLD activation contrasts for the Nback and IdEmo tasks. The global measures were standardized within modality (Z-scores) prior to fitting the GEE model. Since scaled total activation sums to zero, global values for the activation tasks were represented by activation their respective target regions, i.e., midfrontal gyrus for the NBack and averaged activation in amygdala, anterior insula and entorhinal cortex for the IdEmo. While analyzed as continuous variables, age interactions were visually examined by dividing the sample into children (ages less than 13), adolescents (ages 13-17) and young adults (ages 18 and older). Similarly, performance interactions were examined by dividing the sample into high, middle, and low performance bins based on tertile splits of the age-regressed performance scores. These performance splits generated effect sizes by calculating the difference between the top performance tertile and the bottom performance tertile in standard deviation units (Cohen’s D). This method of exploring higher-order interactions by examining not just p-values but also effect sizes was preferred over one that is guided by p-values alone as more reliable and interpretable (Kraemer 2019).

#### 2. Hypothesis-testing

##### Regional analyses: Testing PFIT

To test the PFIT we examined regional specificity and compared regional differences of interactions with performance, contrasting PFIT with non-PFIT regions. Generalized estimating equations (GEE) models were fit within each modality. To minimize type I error given the large number of regions (up to 128 regions per modality), we aggregated them into 8 neuroanatomic Sections: frontal, temporal, parietal, occipital, limbic, baso-striatal, cerebellum and white matter. The PFIT contrast was performed in sections that contained PFIT regions. Section volumes were derived using the sum of all regions involved; for all other brain measures a volume-weighted mean of the regions involved was calculated at the subject-level for each brain section.

For the hypothesis-testing approach, analyses were conducted at the global and regional levels by fitting GEEs with unstructured working correlation structure. GEE models are an extension of generalized linear models that estimate dependence among repeated measures by a user-specified working correlation matrix that allows for correlations in the dependent variable across observations. Five nested forms of GEE models were fit and evaluated as shown below. The null model (Model 1) evaluated the association of the demographic variables, age and sex, on brain parameters. A second model added the performance term to evaluate the association between performance and brain parameters adjusted for demographic variables (Model 2). To evaluate if association of performance and brain parameters differed by sex or by age, interaction terms were added as shown in Model 3 and Model 4 respectively. To evaluate if association of performance and brain differed by both age and sex, Model 5 included all main effects and all possible interactions. Model performance was compared using a Wald test. A squared age term was included to capture non-linear effects of age. Models were fit at all anatomical specificity levels. The hypothesis, PFIT, was tested by contrasting PFIT to non-PFIT regions across the brain with interaction analyses. This analysis was followed by examining the full model, which included brain region as a vector nested within each section.

Model 1: Null model: Sex and Age Associations

Model 2: Model Associations of Performance and Brain

Model 3: Model with Sex modifying the associations of Performance and Brain

Model 4: Model with Age modifying the associations of Performance and Brain

Model 5: Model with Age and Sex modifying the associations of Performance and Brain

For all models above,, indexes the individual brain spatial locations, _*i*_ indicates the participant number, and denotes the random error.

#### 3. Exploratory Hvpothesis-Generation Analyses

The hypothesis-testing phase was followed by an exploratory hypothesis-generation phase, where we used both p-values and effect sizes (see Kraemer 2019) to guide the search for additional regions as candidates for the ExtPFIT network. Exploratory analyses to understand neurodevelopmental changes were conducted for those brain sections and in those modalities that showed significant interactions with performance. Significant effects and interactions of performance were elucidated by charting the brain parameter profiles of effect sizes (Cohen D) for the differences between the high and low performance (tertiles) groups. Regions were rank-ordered for each modality (volume, GMD, etc.) by the p-value of its association with performance and by the effect size separating high from low performance (both after adjusting for covariates including quality metrics and reversing the sign of the effect sizes for MD and CBF, where lower values were hypothesized for the high performance group). The averaged cross-modality ranking was considered evidence of cross-modality relevance to cognitive performance. From each Section containing PFIT regions, we included non-PFIT regions showing a cross-modality ranking comparable or exceeding that of the PFIT region as candidates for inclusion in ExtPFIT. Finally, we examined effect sizes for cerebellar and WM regions for which cross-modality data were available.

#### 4. Data-driven performance-prediction analysis

The CNB-based complex cognition performance factor score (Moore et al. 2015) was predicted using each brain parameter by modality. Subjects included in this analysis passed all modality specific inclusion criteria. Within each modality and then across all modalities, three sets of regressions were fit in training sets, and ***R***^2^s were estimated in the corresponding test sets: 1) performance was regressed on scaled age, age^2^ and age^3^, 2) performance was regressed on scaled brain features subject to a ridge penalty, and 3) performance was regressed on scaled age, age^2^, age^3^, and brain features, with only the brain features subject to the ridge penalty. In each of 10,000 iterations for each set of regressions, the sample was stratified based on performance using the ‘createFolds’ function from the ‘caret’ package (Kuhn et al. 2016) in R into equally-sized training and testing sets. Within each training fold that utilized ridge regression, a model was built using the ‘glmnet’ function in the ‘glmnet’ package (Friedman et al. 2017). The chosen ridge regression penalty parameter minimized out-of-sample mean squared error, where each test set was a unique fifth of the training set for the main ridge regression. The unique variance that brain features can explain in cognition above and beyond age was estimated by taking the difference in the means of the out-of-sample R^2^s from (1) and (3). To test if this difference was significant, differences in ***R***^2^s were calculated for each test set. The proportion of ***R***^2^s in (3) that were greater than (1) for each modality within each sex served as the initial estimates for the p-values, to which FDR correction was then applied to control for multiple comparisons.

## RESULTS

### 1. Global values

The global values showed generally small intercorrelations in either males or females, except for high negative correlations (exceeding .7) between volume and MD and similarly high positive correlations between ALFF and ReHo (Table 1). The small but significant negative correlation between between GMD and CBF, seen both in males and females, confirms that low CBF does not simply reflect low GMD (since CBF is higher in GM than in WM) and is therefore likely of physiological significance. Note that since each modality required specific QA metrics for inclusion, the sample sizes for each modality differed somewhat, and Supplementary Table 1 compares included and excluded groups on sex, age and performance.

The GEE indicated that performance was significantly associated with whole brain global measures across modalities and this association differed by sex and age (GEE, Wald *y2* = 59.51, df=30, p =0.001). This interaction (Figure 1) indicated that while high performers had higher volume and greater GMD compared to medium and low performance groups, they had lower MD and CBF, no differences in ALFF, ReHo or the emotion identification task, and greater activation for the NBack task (first and third rows of Figure 1). Global effect sizes of differences between performance groups became stronger from childhood to adulthood for volume and GMD as well as for NBack activation. For the other parameters this trend seemed more pronounced in males than in females (second and fourth rows in Figure 1).

**Fig. 1.**
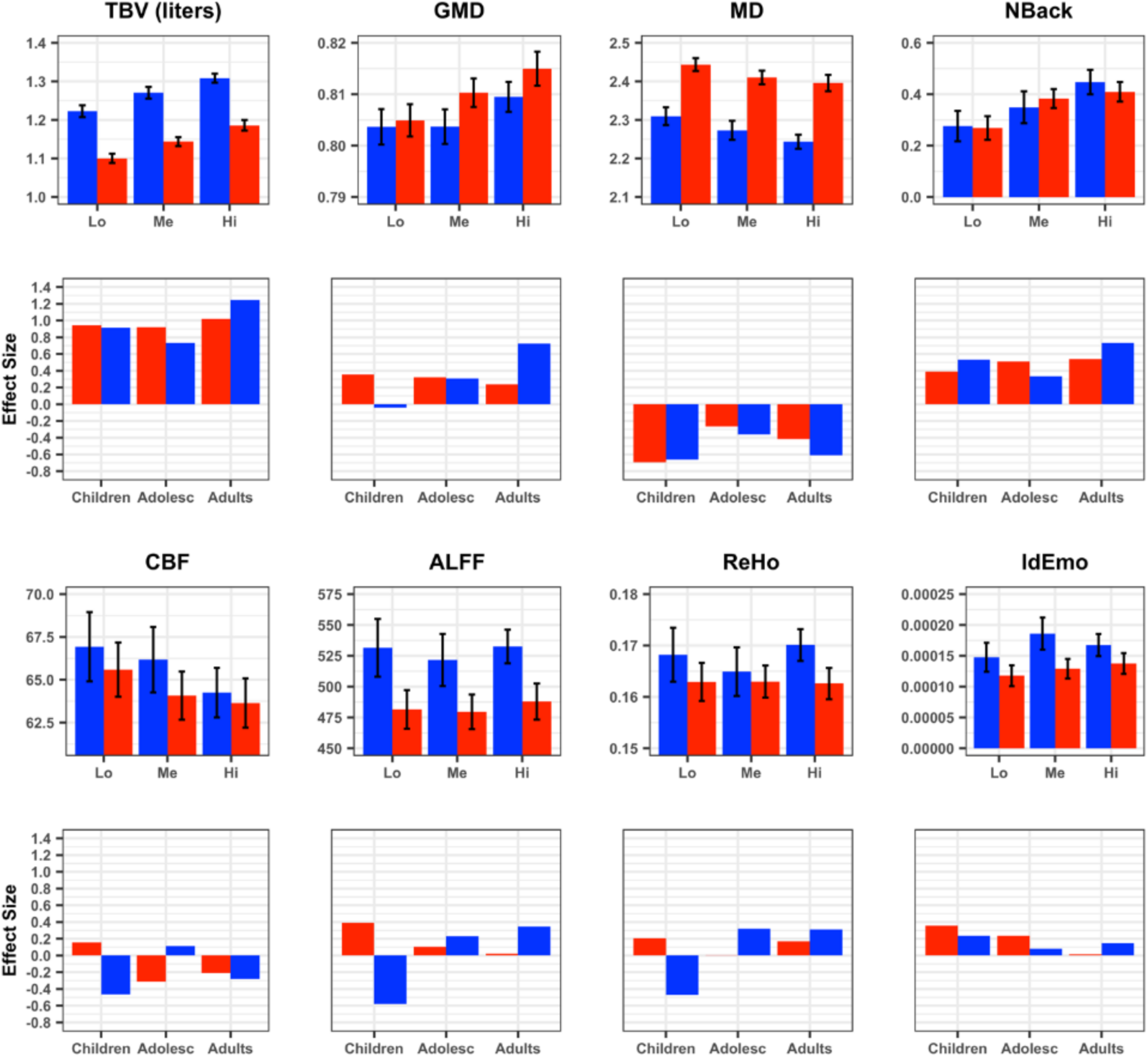
Global values for each modality by performance group (tertiles) in males (blue) and females (red). The first and third panels display means (+95% Cl) of Low (Lo), Medium (Me) and High (Hi) tertiles of performers on Volume, GMD, MD, CBF, ALFF, ReHo, NBack (Medial Frontal Gyrus) and IdEmo (Anterior Insula, Amygdala, Entorhinal Area). The second and fourth panels display effect sizes (Cohen’s D) for the differences between High and Low performers on each parameter in Children, Adolescents and Young Adults.

### 2. Hypothesis testing

The results of the GEE contrasting PFIT to non-PFIT regions are summarized in Table 2. GEEs showed that in all modalities there was a highly significant Performance*PFIT interaction, indicating that performance related to these parameters differently in PFIT and non-PFIT regions. Some higher-order interactions with sex and age were also significant in some modalities, and the 4-way (Performance*PFIT vs non-PFIT*Sex*Age^2^) was significant for GMD. The PFIT regions (Basten, Hilger and Fiebach 2015) and their cross-modality rankings percentiles are illustrated in Figure 2. As can be seen, PFIT regions (blue shaded) generally showed above average or high cross-modality rankings, supporting the PFIT hypothesis.

**Table 2.** Results of the GEE analysis contrasting PFIT to non-PFIT regions with performance, sex, age, age^2^ and the appropriate quality assurance metric as independent vectors.

|  |  | Volume |  | GMD |  | MD |  | CBF |  | ALFF |  | ReHo |  | Nback |  | Idemo |  |
| --- | --- | --- | --- | --- | --- | --- | --- | --- | --- | --- | --- | --- | --- | --- | --- | --- | --- |
| | Df | $\chi^2$ | P | $\chi^2$ | P | $\chi^2$ | P | $\chi^2$ | P | $\chi^2$ | P | $\chi^2$ | P | $\chi^2$ | P | $\chi^2$ | P |
| Perf | 1 | 109.0 | 0.0000 | 135.1 | 0.0000 | 45.6 | 0.0000 | 94.0 | 0.0000 | 4.8 | 0.0292 | 9.8 | 0.0017 | 6.9 | 0.0084 | 2.6 | 0.1069 |
| PFIT | 1 | 119729.0 | 0.0000 | 9037.3 | 0.0000 | 0.3 | 0.6005 | 14853.0 | 0.0000 | 4617.2 | 0.0000 | 7887.2 | 0.0000 | 1235.1 | 0.0000 | 3122.7 | 0.0000 |
| Sex | 1 | 547.0 | 0.0000 | 17.8 | 0.0000 | 349.8 | 0.0000 | 7.0 | 0.0106 | 43.9 | 0.0000 | 11.1 | 0.0009 | 0.1 | 0.7307 | 5.6 | 0.0182 |
| Age | 1 | 63.0 | 0.0000 | 211.1 | 0.0000 | 12.2 | 0.0005 | 141.0 | 0.0000 | 87.8 | 0.0000 | 153.2 | 0.0000 | 0.1 | 0.7956 | 0.3 | 0.6182 |
| Age2 | 1 | 1.0 | 0.3902 | 0.7 | 0.4065 | 7.7 | 0.0055 | 32.0 | 0.0000 | 0.8 | 0.3585 | 0.4 | 0.5137 | 0.1 | 0.7154 | 0.1 | 0.7061 |
| QA | 1 | 1.0 | 0.4591 | 235.1 | 0.0000 | 4.6 | 0.0324 | 2.0 | 0.1980 | 30 | 0.0000 | 30.9 | 0.0000 | 0.2 | 0.7001 | 4.3 | 0.0389 |
| Perf*PFIT | 1 | **235.0** | **0.0000** | **5.7** | **0.0172** | **12.3** | **0.0004** | **44.0** | **0.0000** | **26.7** | **0.0000** | **24.6** | **0.0000** | **47.8** | **0.0000** | **33.5** | **0.0000** |
| Perf*Sex | 1 | 1.0 | 0.2671 | 2.2 | 0.1394 | 5.9 | 0.0149 | 11.0 | 0.0009 | 0.2 | 0.6321 | 0.6 | 0.4432 | 0.8 | 0.3877 | 0.3 | 0.5640 |
| PFIT*Sex | 1 | 465.0 | 0.0000 | 105.7 | 0.0000 | 279.0 | 0.0000 | 5.0 | 0.0328 | 0.7 | 0.3903 | 5.7 | 0.0166 | 5.9 | 0.0155 | 0.3 | 0.5824 |
| Perf*Age | 1 | 4.0 | 0.0520 | 4.1 | 0.0430 | 0.1 | 0.7183 | 1.0 | 0.2236 | 0.1 | 0.7336 | 0.6 | 0.4405 | 0.1 | 0.8275 | 0.3 | 0.5724 |
| Perf*Age2 | 1 | 0.0 | 0.5722 | 0.1 | 0.7103 | 0.1 | 0.8271 | 0.0 | 0.5546 | 1 | 0.3094 | 0.1 | 0.8194 | 2.7 | 0.1036 | 0.2 | 0.6596 |
| PFIT*Age | 1 | 1.0 | 0.2283 | 2.7 | 0.1002 | 76.5 | 0.0000 | 88.0 | 0.0000 | 7.1 | 0.0079 | 2 | 0.1600 | 3.3 | 0.0714 | 17.5 | 0.0000 |
| PFIT*Age2 | 1 | 0.0 | 0.6158 | 3.8 | 0.0498 | 1.4 | 0.2349 | 3.0 | 0.0631 | 11.4 | 0.0007 | 0.4 | 0.5261 | 3.8 | 0.0522 | 0.5 | 0.4708 |
| Sex*Age | 1 | 8.0 | 0.0060 | 0.6 | 0.4386 | 10.2 | 0.0014 | 25.0 | 0.0000 | 0.1 | 0.7079 | 0.6 | 0.4357 | 0.0 | 0.8594 | 0.0 | 0.8325 |
| Sex*Age2 | 1 | 8.0 | 0.0047 | 0.1 | 0.8067 | 2.0 | 0.1605 | 5.0 | 0.0210 | 0.2 | 0.6488 | 1.4 | 0.2310 | 1.5 | 0.2166 | 1.4 | 0.2426 |
| Perf*PFIT*Sex | 1 | 1.0 | 0.2788 | 3.4 | 0.0655 | **8.5** | **0.0035** | **8.0** | **0.0037** | 2.4 | 0.1198 | 0.6 | 0.4293 | 0.0 | 0.9731 | 2.3 | 0.1280 |
| Perf*PFIT*Age | 1 | **5.0** | **0.0233** | **10.3** | **0.0013** | 1.2 | 0.2730 | 0.0 | 0.7808 | 0.3 | 0.5755 | 0.4 | 0.5144 | 0.0 | 0.8422 | 0.3 | 0.6045 |
| Perf*PFIT*Age2 | 1 | 0.0 | 0.8277 | 0.7 | 0.4171 | 0.5 | 0.4924 | 0.0 | 0.7909 | 0.7 | 0.4003 | 0.1 | 0.7038 | **6.1** | **0.0133** | 0.0 | 0.8360 |
| Perf*Sex*Age | 1 | 1.0 | 0.2730 | 0.4 | 0.5266 | 4.2 | 0.0396 | 0.0 | 0.4874 | **8.6** | **0.0034** | 3.5 | 0.0630 | 0.0 | 0.8607 | 0.2 | 0.6797 |
| Perf*Sex*Age2 | 1 | 1.0 | 0.2310 | 0.5 | 0.4883 | 0.0 | 0.9979 | 3.0 | 0.1134 | 2.8 | 0.0942 | 1.6 | 0.2037 | 0.7 | 0.4164 | 1.3 | 0.2624 |
| PFIT*Sex*Age | 1 | 6.0 | 0.0149 | 7.0 | 0.0081 | 0.9 | 0.3549 | 11.0 | 0.0010 | 0.5 | 0.4995 | 0.3 | 0.5795 | 0.1 | 0.8055 | 5.9 | 0.0150 |
| PFIT*Sex*Age2 | 1 | 8.0 | 0.0044 | 13.0 | 0.0003 | 1.7 | 0.1868 | 0.0 | 0.7436 | 0 | 0.8558 | 0 | 0.9393 | 0.4 | 0.5480 | 0.3 | 0.5795 |
| Perf*PFIT*Sex*Age | 1 | 1.0 | 0.3340 | 2.0 | 0.1598 | 2.3 | 0.1311 | 0.0 | 0.5492 | 1.6 | 0.2088 | 0 | 0.8712 | 0.0 | 0.8808 | 4.1 | 0.0440 |
| Perf*PFIT*Sex*Age2 | 1 | 1.0 | 0.3957 | **7.5** | **0.0061** | 1.5 | 0.2226 | 0.0 | 0.5524 | 0.2 | 0.6527 | 0.3 | 0.5629 | 0.1 | 0.8123 | 0.1 | 0.7339 |

Perf=Performance score; PFIT=Parieto-Frontal Integration Theory regions; QA=Quality assurance metric
**Bold** = Significant; **Bold Italics** =Interacts with Performance; *Italics* =Marginally significant interaction with performance

**Fig 2.**
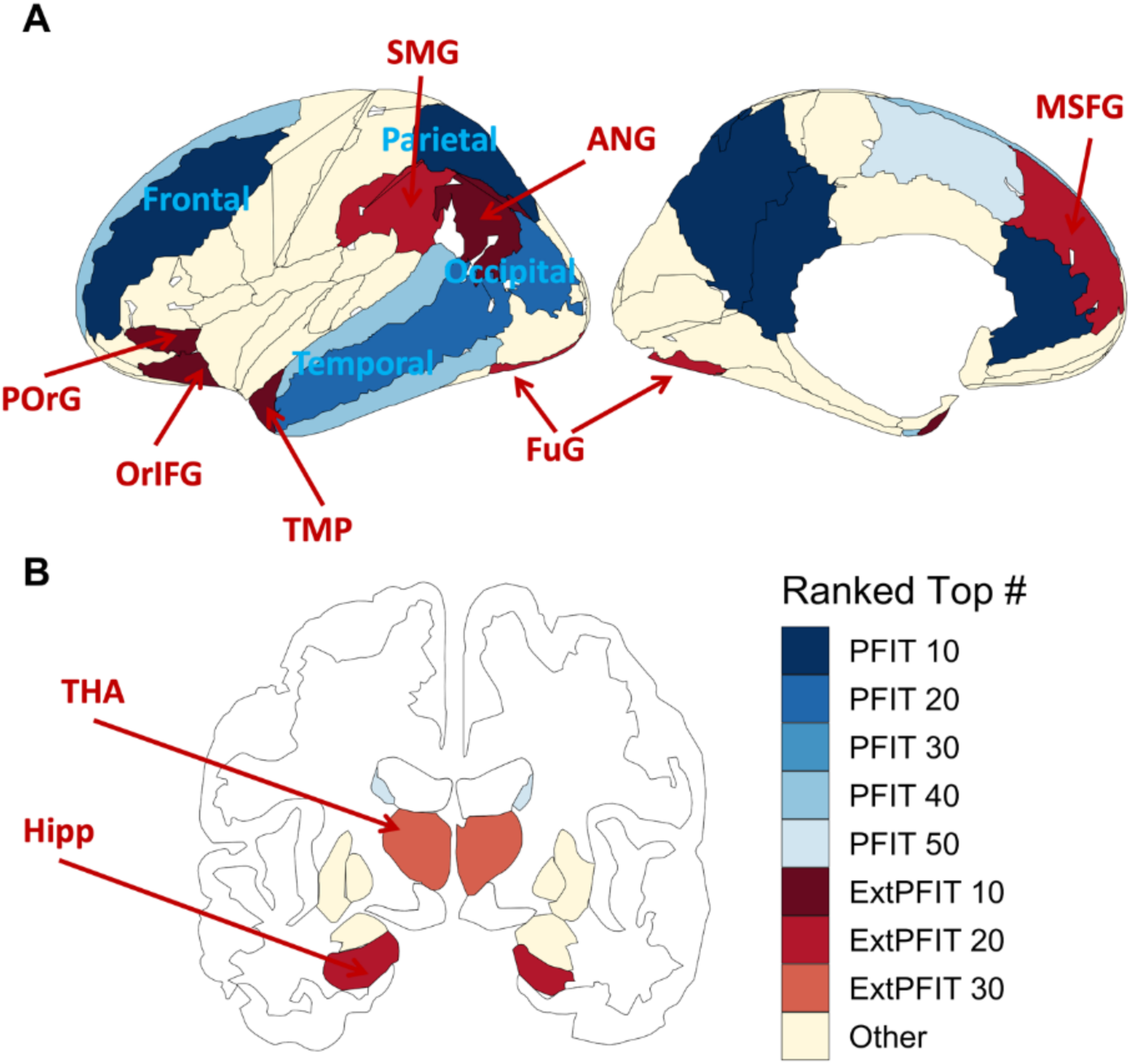
The image displays the number of modalities that ranked at the top when averaging p-value and effect sizes, colored by PFIT status. PFIT regions are in shades of blue, and extended-PFIT (ExtPFIT) regions are in shades of red. The p-values and effect sizes are for the performance predicting the normalized brain feature, controlling for normalized quality, age, age squared and age cubed. Quality metrics are as follows: volume and gray matter density use the average manual rating; mean diffusivity uses the temporal signal to noise ratio; and cerebral blood flow, amplitude of low frequency fluctuations, regional homogeneity, NBack and IdEmo use the mean of the relative root mean square displacements.

### 3. Exploratory analyses

The exploratory GEE model did not contrast PFIT to non-PFIT regions, but instead included Region as a within-group (“repeated-measures”) vector nested within each of the 8 brain sections. All other effects (Performance, Sex, Age, Age^2^ and QA metric) were the same, and we tested for all Performance interactions in each of the 8 Sections (Supplementary Table S2). The results showed highly significant (all p<.001) Performance*Region interactions in each and every modality for frontal, temporal and parietal sections, and in all sections for volume, GMD and NBack. These interactions indicate regionally-specific relations to performance in each brain section. Higher-order interactions involving age and sex were absent for volume but were highly significant in the other modalities, indicating that for brain parameters other than volume the relation between performance and brain parameters shows developmentally related sex differences.

To characterize the involvement of different parameters related to cognitive performance, we examined the cross-modality rankings in non-PFIT regions to identify candidates for inclusion in an ExtPFIT network - after controlling for age, sex and scan quality (including motion). As can be seen in Figure 2, several regions were strong candidates for ExtPFIT (brown shaded) by showing performance-related cross modality rankings that are comparable to those seen in PFIT regions. These results suggest that the PFIT should consider expansion to incorporate additional frontal (orbital, inferior and precentral, and mid superior frontal gyrus), temporal (temporal pole) parietal (supramarginal and angular gyri) and occipito-temporal cortex (lingual and fusiform), as well as limbic (hippocampus) and baso-striatal (thalamus) components.

To examine how these associations between brain parameters and performance are manifested in each modality and how they develop in males and females, we plotted effect sizes for PFIT and ExtPFIT regions. These results are detailed below.

#### a. Neuroanatomic parameters

The regional distribution of the PFIT and ExtPFIT effect sizes, contrasting high and low performers in children, adolescents and young adults on each of the neuroanatomic parameters are shown in Figure 3.

**Fig. 3.**
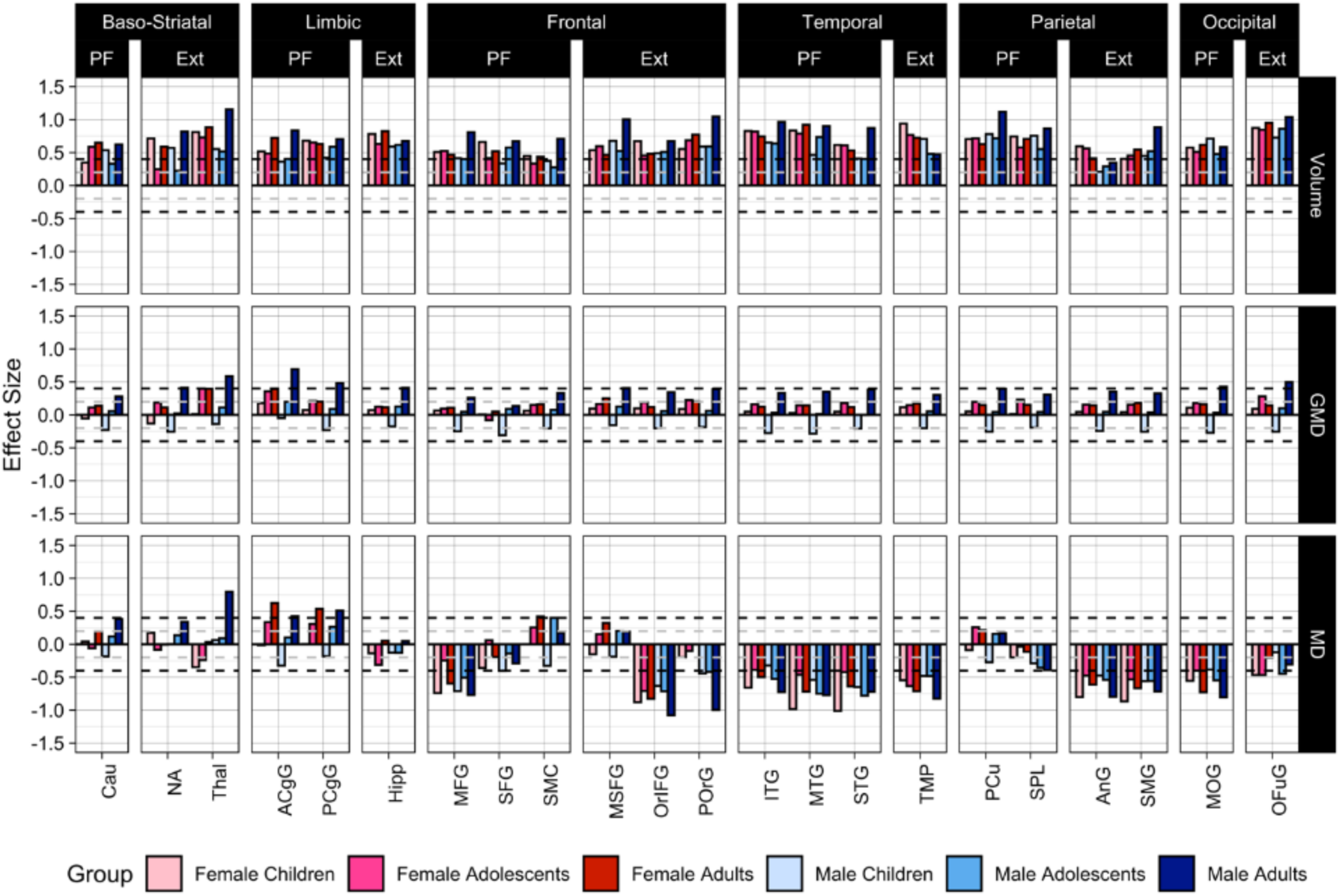
Developmental and sex effects for the association between structural brain features and high versus low performers are displayed. Effects shown are for volume (upper panel), gray matter density (GMD; middle panel) and mean diffusivity (MD; lower panel). Females are in shades of red from light (children) to pink (adolescents) to red (adults) while males are similarly displayed in shades of blue. Effect sizes are the coefficients for the indicator variable for the high compared to low performers predicting the scaled brain feature, controlling for scaled quality, age, age squared and age cubed within each age bin. The quality metric for volume and gray matter density is the average manual rating, and for mean diffusivity is the temporal signal to noise ratio. The subset of non-PFIT regions that rank at or above PFIT regions, and therefore are considered for the ExtPFIT, are also displayed. Horizontal dashed lines indicate the transition from small to moderate effect sizes. Regional abbreviations: Cau=Caudate nucleus; NA=Nucleus Accumbens; Thal=Thalamus; ACgG=Anterior Cingulate Gyrus; PCgG=Posterior Cingulate Gyrus; Hipp=Hippocampus; MFG=Middle Frontal Gyrus; SFG=Superior Frontal Gyrus; SMC=Supplementary Motor Cortex; MSFG=Superior Frontal Gyrus Medial Segment; OrIFG=Orbital part of the Inferior Frontal Gyrus; POrG=Posterior Orbital Gyrus; ITG=Inferior Temporal Gyrus; MTG=Middle Temporal Gyrus; STG=Superior Temporal Gyrus; TMP=Temporal Pole; PCu=Precuneus; SPL=Superior Parietal Lobule; AnG=Angular Gyrus; SMG=Supramarginal Gyrus; MOG=Middle Occipital Gyrus; OFuG=Occipital Fusiform Gyrus.

As can be seen in Figure 3, for volume (upper row) all PFIT regions show moderate to large effect sizes indicating higher volumes in the high-performance groups across age groups and in both males and females. This effect increased linearly with advanced age in all regions for males, although there are several PFIT regions where children and adults have larger effect sizes than adolescents, while in females the effect reached adult size already during childhood in some regions. Several ExtPFIT regions showed comparable effect sizes (Figure 3, upper row, regions under ‘Ext’), and they include the nucleus accumbens and thalamus, implicating reward and relay system components in complex cognition. The hippocampus is a limbic region that showed similar effect sizes as the anterior and posterior cingulate, the only limbic regions thus far implicated by PFIT (Basten, Hilger and Fiebach 2015). Similarly, superior, midfrontal and somatomotor cortex were the only frontal regions included in PFIT, while our results indicate equal or higher effect sizes for the medial superior frontal gyrus and orbital (both inferior and posterior) cortex. Of the temporal lobe regions, in addition to inferior, superior and midtemporal regions, the temporal pole showed robust effect sizes, and for parietal regions the angular and supramarginal gyri showed similar effect sizes to the PFIT precuneus and the superior parietal lobule. Finally, the occipital fusiform cortex showed effect sizes similar and even exceeding the PFIT mid-occipital region.

For GMD we observed considerably smaller effect sizes compared to volume (Figure 3, middle row). For PFIT relative to non-PFIT regions this association differed by sex and by age^2^ (Table 2), and this was the case for the frontal, parietal and occipital lobes (Supplementary Table 2). High performers had higher GMD in PFIT regions across age and sex groups, except for male children, and in males the effect sizes increased in most regions from childhood (negative) to adolescence to young adulthood (positive), while in females they remained generally stable across age bins. ExtPFIT regions (Figure 3, middle row, regions under ‘ExtPFIT’) likewise showed small to moderate effect sizes in the direction of higher values in the high-performance group, with largest effects in the adult males. The ExtPFIT regions did not differ in performance group effects from the original PFIT regions.

MD showed much larger effect sizes than GMD in the direction of lower values in high performers compared to low performers (Figure 3, lower row). Indeed, these effect sizes approached those for volume in frontal, temporal and occipital PFIT regions, although not in parietal PFIT regions. These effects appeared across age groups but became most pronounced with advanced developmental age, especially in males. For the parietal component of PFIT this effect was seen only in the superior parietal lobule and not in precuneus, and the effects on limbic components of PFIT were in the opposite direction. Several ExtPFIT regions, specifically frontal and parietal, showed similar effect sizes in the same direction, and the effect is evident in ExtPFIT parietal regions, where it was absent in PFIT parietal regions.

#### b. Neurophysiologic parameters

Effect sizes for the neurophysiologic parameters are shown in Figure 4. For CBF (Figure 4, top row), PFIT regions showed small to moderate effect sizes, some reaching -.5 SDs, of lower CBF associated with high performance. These associations were stronger for males than females and showed bigger increase from adolescence to young adulthood in males. Several ExtPFIT regions showed similar effect sizes in the same direction, including baso-striatal (nucleus accumbens and thalamus) and limbic (hippocampus) regions.

**Fig. 4.**
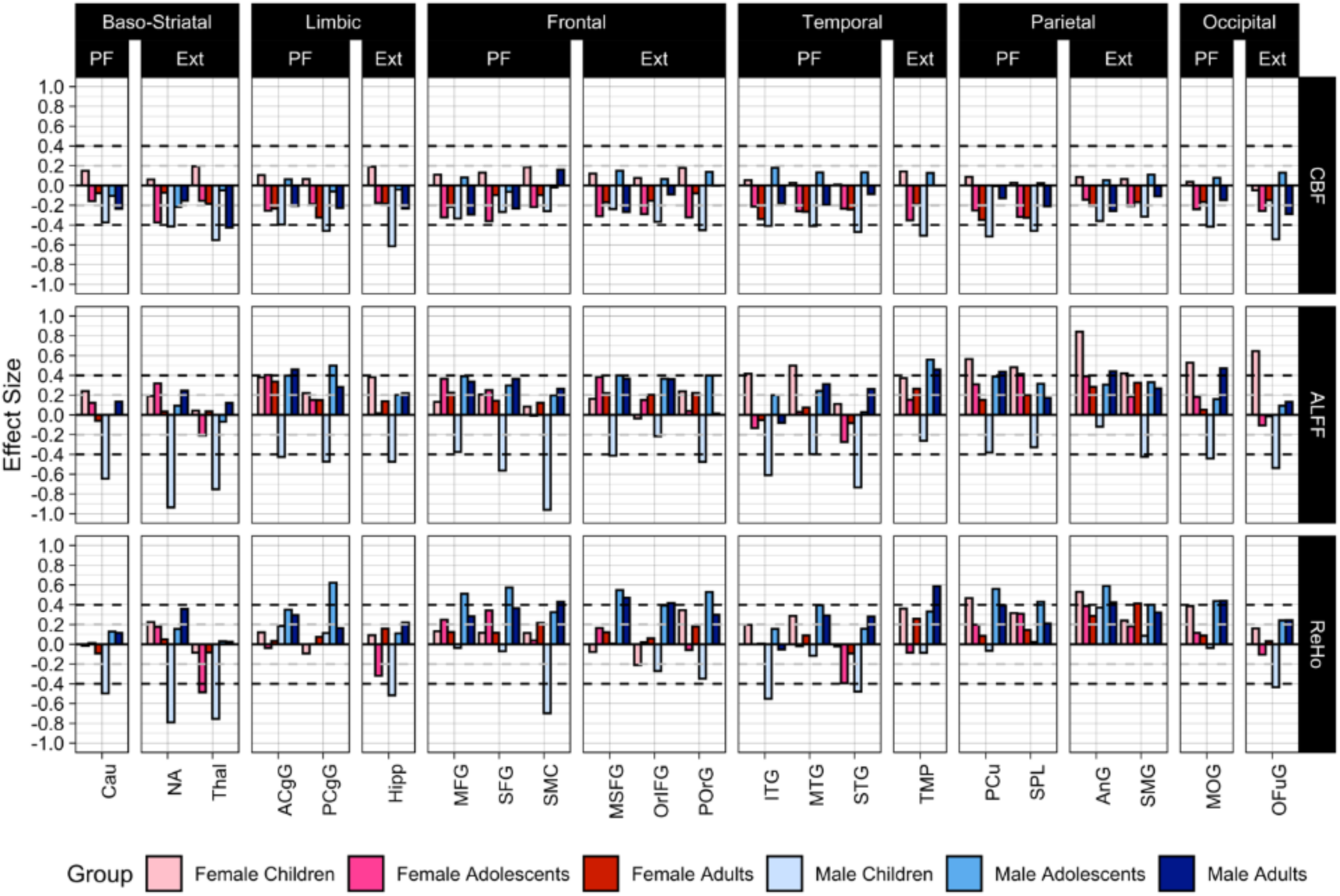
Developmental and sex effects for the association between structural brain features and high versus low performers are displayed. The quality metric for cerebral blood flow, amplitude of low frequency fluctuations, and regional homogeneity is the mean of the relative root mean square displacements. All other values, legends and abbreviations are as in Figure 3.

For ALFF (Figure 4, middle row), high performers had higher values in most PFIT regions and across age groups with small to moderate effect sizes (.2 to .5 SDs), with the exception of male children who showed moderate to large effect sizes (up to -.9 SDs) in the opposite direction. Similar effects were seen in most ExtPFIT regions. Effect sizes for ReHo (Figure 4, bottom row) were of similar direction and magnitude to those for ALFF.

Effect sizes for the activated fMRI are presented in Figure 5. The results for the working memory (NBack) task (upper row) give remarkably strong support for the narrow PFIT model, as moderate to large effect sizes are seen across the age groups for frontal and parietal regions from the original PFIT. Notably, the thalamus, from the ExtPFIT, shows a similar effect size, supporting its role in cognition. In contrast to the NBack, effect sizes are generally negligible to small for the emotion identification task (Figure 5, lower row).

**Fig. 5.**
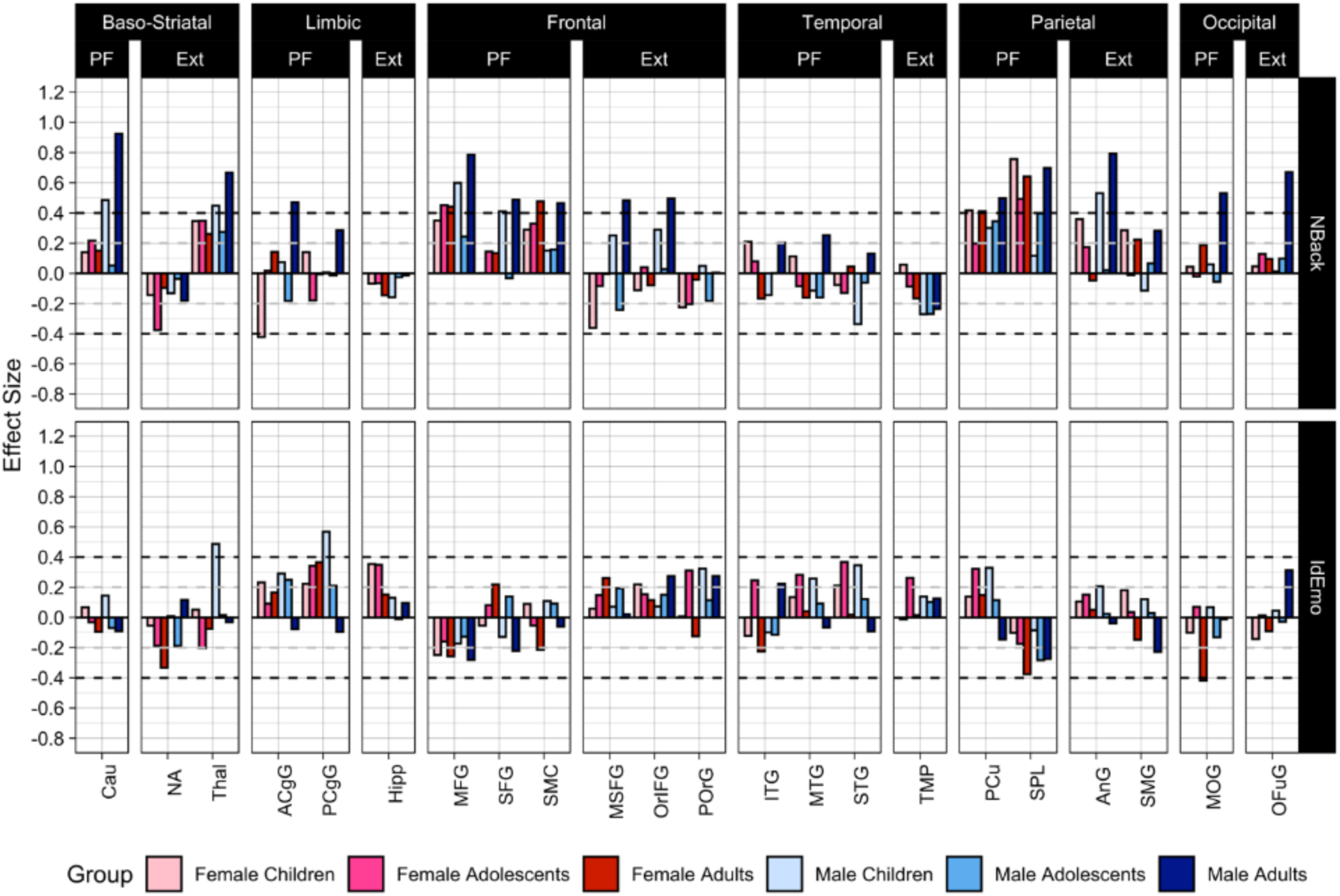
Developmental and sex effects for the association between basal performance and activation on the NBack and IdEmo tasks in the same regions as in previous Figures. The quality metric for these effects is the mean of the relative root mean square displacements. All other values, legends and abbreviations are as in Figure 3.

##### Cerebellum and white matter

As can be seen in Figure 6, moderate to large effect sizes separating high and low performers were seen for volume in some cerebellar regions and all WM regions. These effect sizes were seen in all age groups but were most pronounced in the oldest group of young adult males, where they reached and sometimes exceeded 1SD. For WM, MD and CBF showed small to moderate effects in the direction of lower values associated with better performance. The cerebellar regions showing the most robust effect sizes were lobules 8-10 and Exterior cerebellum. Effect sizes in other modalities measured were small to moderate; they are shown for future reference and will not be further discussed.

**Fig. 6.**
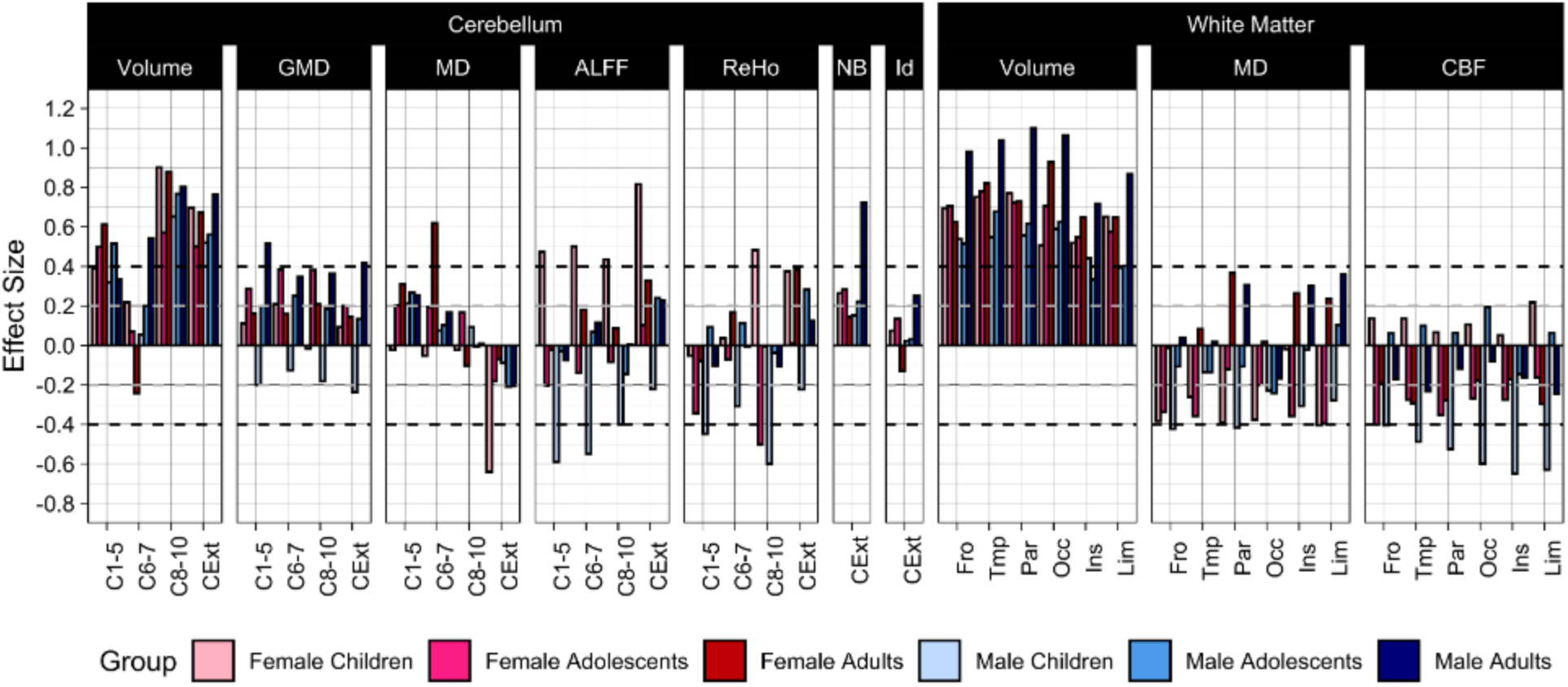
Developmental and sex effects for the association between basal performance and cerebellar and white matter features comparing high versus low performers. Quality metrics are as described for each modality in the above captions. Abbreviations: Cl-5=Cerebellar layers I to V; C6-7=Cerebellar layers VI to VII; C8-10=Cerebellar layers VIII to X; CExt=Cerebellar exterior; Fro=Frontal; Temp=Temporal; Par=Parietal; Occ=Occipital; Ins=Insular; Lim=Limbic. Other values and legends are as in Figure 3.

### 4. Data-driven analyses

Each modality considered alone explained at least 9% of the variance in performance, estimated out-of-sample, with volume explaining the greatest proportion of the variance for both females and males (31.9 and 32.1%, respectively). However, since in this age range of 8 to 22 years both performance and brain parameters are highly correlated with age, we aimed to establish how much variance in performance is explained by brain parameters above and beyond age. We found that volume, gray matter density, NBack activation and all of the modalities combined explained a significant amount of variance in cognition above and beyond age for females, while only volume did so for males (Figure 7).

**Fig 7.**
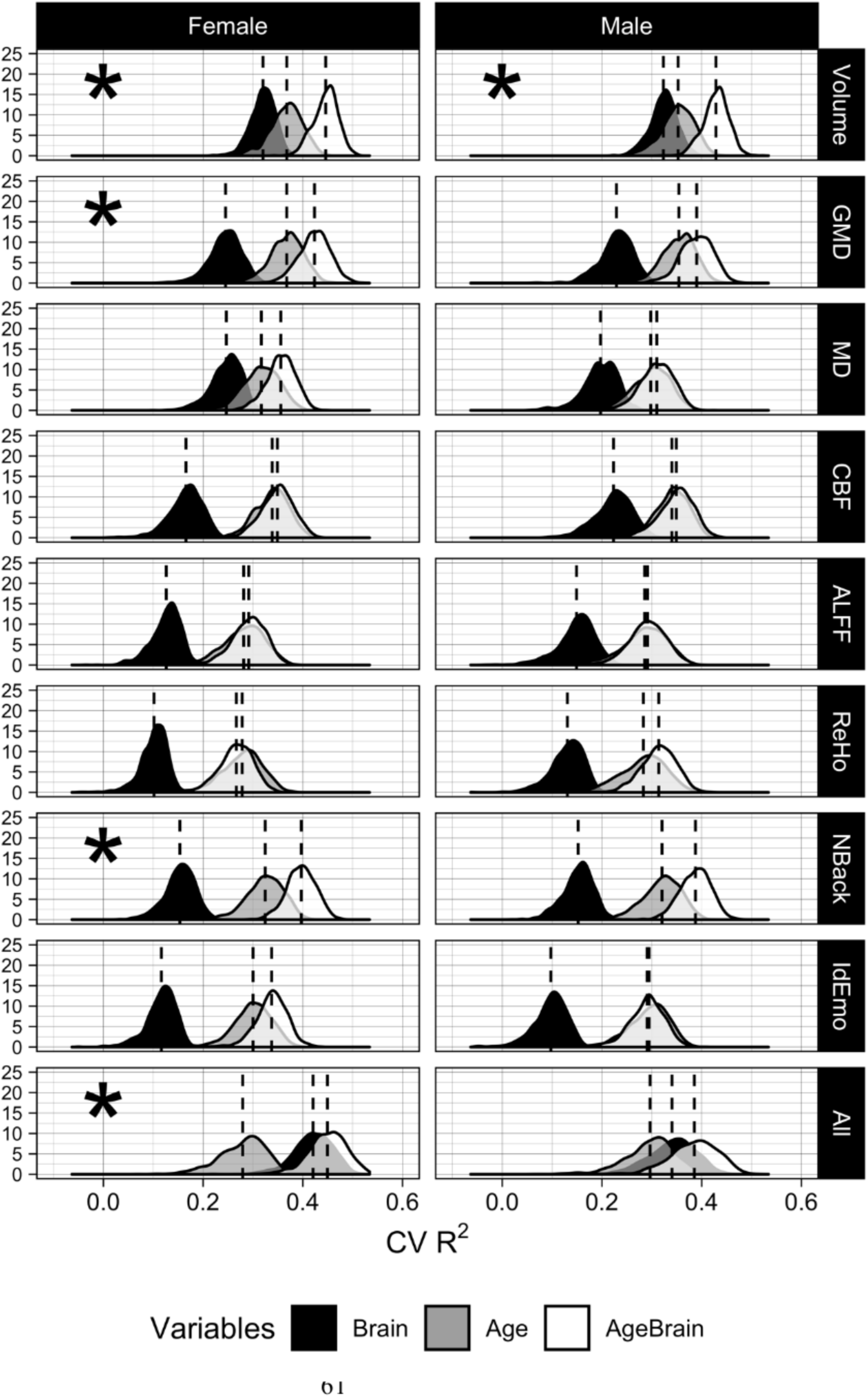
Results of ridge regressions predicting the executive functioning and complex cognition factor score for each modality and using all modalities (“All”). The distributions shown are out- of-sample R^2^s over 10,000 iterations. For each R^2^, a random half of the data was assigned to the training set, and the other half was assigned to the test set. R^2^ was calculated using the following formula:. The ridge hyperparameter lambda was chosen using five-fold cross-validation on the training data. The lambda that minimized the out-of-sample MSE was selected.

## DISCUSSION

Our results offer fresh insights regarding how cognition is related to multimodal parameters of brain structure and function. Global values for anatomic and physiologic parameters were associated with performance in modality-specific ways. Overall, high cognitive performers had anatomically higher volumes, higher GMD and lower MD, and physiologically they evinced lower CBF and higher activation to the NBack task. In all those modalities, the age effects were significant and effect sizes separating high from low performers generally increased with age. Performance associations with global ALFF and ReHo were more complex and differed between males and females. Regional analyses (see below) indicated more consistent effects in specific regions. The results of the global values indicate that the brain parameters examined relate meaningfully to performance and that this relationship is strengthened during development.

The theoretical thrust of the analysis was to examine whether anatomic and physiologic indices derived from the network of regions included in PFIT indeed show association with cognitive performance, and whether other regions show similar performance associations warranting their inclusion in an ExtPFIT network. We found strong support for cross-modality involvement of PFIT regions, which showed significantly greater association with cognitive performance than non-PFIT regions in all modalities (see Table 2), and these relations were modulated only by age for volume and by higher order interactions of age and sex for other parameters. Examination of effect sizes contrasting high and low performers indicated that they were large (up to > 1 SDs for adult males, see Figure 3 top row) for higher volume and moderate for higher GMD, lower MD, lower CBF, higher ALFF and ReHo, and greater activation to the NBack task. The amplitude and direction of results in this multimodal study are consistent with the literature and meta-analyses where subsets of these parameters have been examined in specific studies (Ritchie et al. 2015, Ryman et al. 2016, Gen<; et al. 2018, Yan et al. 2011, Hshieh et al. 2017, Rane et al. 2018, Pamplona et al. 2015, Vakhtin et al. 2014, Finn et al. 2015, Yoo et al. 2018, Fong et al. 2019, Greene, Gao, Scheinost, & Constable 2018, Dubois, Galdi, Paul and Adolphs 2018).

However, several regions not included in PFIT showed comparable cross-modality effect sizes and merited consideration for inclusion in an extended PFIT. These regions included some that surround the original PFIT regions, indicating that larger portions of frontal, temporal, parietal and occipital cortex are involved in optimizing complex cognition. The frontal regions that should be considered for inclusion in ExtPFIT are adjacent to the PFIT but more posterior (somatosensory gyrus), medial (medial superior frontal gyrus), and inferior (orbital), suggesting the contribution of top-down regulatory systems for cognitive performance (Hampshire et al. 2010, Swick, Ashley and Turken 2008, Rolls and Grabenhorst 2008, Wojtasik et al. 2020). The parietal candidates for ExtPFIT are the supramarginal and angular gyri, regions long implicated in complex cognition (Geschwind 1970, Tremblay and Dick 2016), as is the temporal pole, a region implicated in multimodal sensory integration (Olson, Plotzker and Ezzyat, 2007) and social cognition (Pehrs et al. 2017), underscoring the role of temporal lobe connectivity in complex cognition (Blazquez Freches et al. 2020). The occipital region that could be included in ExtPFIT, fusiform gyrus, is involved in high-order visual processing and would contribute visual memory and concept formation (Mechelli et al. 2000). In addition to these cortical areas, baso-striatal and limbic regions should be considered for ExtPFIT in addition to the caudate and cingulate included in the PFIT. The ExtPFIT seems to include the accumbens and thalamus, reinforcing the striatal component that implicates motivational aspects of cognitive performance. Indeed, subcortical regions have been implicated in higher-order cognitive function and memory (Koziol et al. 2014; Miinte et el. 2008; Wolff and Vann 2019), and ventral striatum activation signaling internal reward during NBack correlated with performance in the PNC (Satterthwaite et al. 2012). The limbic region added to the ExtPFIT model is the hippocampus, implicating contributions of emotion regulation and episodic and contextual memory integration (Puigdemont et al. 2012, Aminoff, Kveraga and Bar 2013). Thus, the original PFIT emphasized frontal and parietal regions and subsequent meta-analyses implicated some additional temporal, occipital, limbic and striatal regions as important nodes of the complex cognition network. Our results from a multimodal study of a single large sample indicate that the network should be broadened, and optimal cognitive performance relates to a multimodal network with robust representation of regions required for integrating conceptual processing with perception, memory, emotion regulation, and motivation.

Additional regions that show robust effect sizes related to performance were found in the cerebellum, specifically exterior and lobules 8-10. The cerebellum, traditionally considered primarily in relation to motor function and coordination, has been increasingly recognized as a hub related to cognition (Koziol et al. 2014) and these regions within the cerebellum have been specifically implicated in cognition (Tedesco et al. 2011). In addition, all white matter regions showed sizable effect sizes, ranging from .4 to >1 SD, with better performance associated with higher volumes, as well as reduced MD and CBF. These findings indicate that white matter integrity and tuning contribute to optimize cognitive performance.

The sample’s age range from childhood to young adulthood allowed examination of developmental effects and sex differences in the magnitude of effect sizes related to performance. For volume, these effect sizes were stable across age groups, with a significant interaction with age showing that they tend to increase in size from childhood to adulthood. For the other modalities, the results showed higher order interactions with age and sex, indicating that these parameters become optimized for performance over development at different rates in males and females. The sex differences overall seem to indicate complementary mechanisms in males and females that serve to compensate for sex differences across brain parameters. Sex differences in brain-behavior associations have been well documented (e.g., Faraone and Tsuang 2001, Gur et al. 1982, 1999, 2000; Jazin and Cahill 2010, Ragland et al. 2000, Raznahan et al. 2011, Satterthwaite et al. 2015). Thus, lower volume in females is compensated for by higher GMD, and performance is further modulated by MD, CBF, ALFF and ReHo, which may together account for equal cognitive performance. Furthermore, age-related differences in this developmental cohort were smaller for females than for males, indicating that females reach adult differences earlier, consistent with other studies (Erus et al. 2014; Goyal et al. 2019). We might speculate that earlier stabilization of metabolic parameters in females helps sustain brain integrity throughout the adult lifespan. Such complementarity between the sexes might have enhanced survival and reproduction in humans’ environment of evolutionary adaptedness (Barkow, Cosmides, and Tooby 1995). These sex differences could further reflect complementary reliance on different aspects of brain structure and function to optimize cognitive performance.

The data-driven analysis indicated that volume was far and away the brain parameter most strongly associated with cognitive performance, confirming earlier findings associating higher brain volumes with better cognitive abilities (Witelson, Beresh, and Kigar 2005; Gignac and Bates, 2017; Nave et al. 2018; Pietschnig et al. 2015). In our sample, effect sizes separating high from low performance groups were moderate to large, and cross-validated R^2^s for predicting performance based on volume alone exceeded .3 for both females (32.01% of variance) and males (32.33%) (see Figure 7). This estimate of explained variance is at the upper-range of estimates from prior studies, which vary from 3% to >30%. Most previous studies to which we can compare our results examined volume. Our R^2^ values are considerably higher than those reported for volumes by Nave et al. (2018), who estimate that volume explains slightly over 3% of the variance in cognitive performance in an adult sample (age range 40 to 69 years). Possibly the more extensive battery on which our performance measure was based, as well as the use of the same scanner, could have eliminated some sources of noise in estimating the dependent measures. We also established that high GMD relates to better performance, although the variance explained beyond age was more modest (5.58% of variance for females and 3.56% for males). The other parameters showed smaller predictive power, except for the activation to the NBack task, which uniquely explained substantial amount of out-of-scanner performance. These results are consistent with earlier work showing that task-activated fMRI is a better predictor of performance than resting-state measures (Greene, Gao, Scheinost, and Constable 2018; Yoo et al. 2018). Overall, considering the inherent error in all our measurements, these results indicate substantial coupling between cognitive performance and parameters of brain structure and function.

It is notable that CBF was measured at a resting state, characterized as the “default mode” (Raichle et al., 2001) condition. Our finding that lower resting-state CBF is associated with better overall performance is consistent with reports that deactivation of the default-mode network during task performance is as predictive of performance as activation of task-related regions (Satterthwaite et al., 2013). A lower basal metabolic rate, as indicated by lower basal CBF, could be indicative of greater metabolic efficiency and potentially suggest a greater dynamic range in brain function. Thus, a lower “idling rate” may be conducive to better performance by permitting activation when the individual is faced with a task while preserving energy in the absence of a task.

Several limitations of this study are noteworthy. First and foremost, the study is cross-sectional and therefore unable to evaluate developmental trajectories of performance and brain parameters. All conclusions regarding age-related differences are limited by this feature of the data and longitudinal studies are needed to establish trajectories of the observed associations. The age range of the sample, 8 to 22 years, limits generalizability to other ages. Within this age range, in which age related differences were seen in all brain modalities examined, the performance-related differences were generally consistent and replicated in all age groups. Another limitation of the study was the focus on a single parameter of cognitive capacity. This focus was necessitated by the complexity of probing multiple brain regions across modalities and accounting for age effects and sex differences. The measure selected is the closest proxy for “IQ”, which was used in other studies and thus improves comparability of results. Future analyses can focus on other performance domains, such as episodic memory and social cognition, and more specific aspects of performance such as accuracy compared to speed. The study is also limited by analyzing data across the entire sample, which is quite heterogeneous and, while ascertained through general pediatric services and not psychiatric services, still included individuals with significant psychopathology (Calkins et al. 2015) and adverse life events (Barzilay et al. 2018, 2020), and from diverse sociodemographic, ethnic and racial backgrounds. Perhaps stronger and more coherent effects could be seen if we limited the analyses to the subsample of typically developing youth without any significant disorder or to more homogeneous populations. We believe that while such analyses have merit and could reveal effects of different disorders on the observed relationships, the heterogeneity and diversity of our sample enhance generalizability of the reported results. Additionally, our analyses examined regional parameters of brain structure and function and related them to individual differences in behavioral measures of cognitive performance taken within the same timeframe, but not contemporaneously in the scanner. More variance in behavioral measures could be linked to functional brain parameters acquired contemporaneously (Roalf et al. 2014). Finally, our analyses are limited by variability among parameters in the reliability of measures and by the high dimensionality of the data necessitating control for multiple comparisons. The reliability of the measures used in this study is acceptable to high, and we have incorporated approaches to reduce data dimensionality and contain Type I error, but our approach may have obscured important findings that future work, better addressing these issues, can reveal.

Notwithstanding its limitations, the present study provides some “benchmarks” for assessing relations among brain parameters and performance. The results can guide hypotheses on how brain structure and function relate to individual differences in cognitive capacity, and offer the ability to gauge the relevance to cognitive performance of group differences or changes in brain parameters. Furthermore, acquisition of each parameter is costly in time and data management resources, and our study can inform design of future large-scale neuroimaging studies based on the relevance of associating acquired brain parameters with cognitive performance. Since volume and GMD measures can be obtained rapidly and have shown the least susceptibility to QA failure in our data, they have key advantages in studies seeking to establish neural substrates of cognition. Activated fMRI can offer more specific associations to performance than resting-state physiologic measures, and multimodal collections including future efforts at multivariate integration across modalities could take advantage of their complementary strengths and weaknesses. Our finding of lower resting CBF in high performers is worthy of special emphasis since, unlike the structural parameters of volume, GMD and MD, it relates to brain function. Uncovering a physiologic index associated with individual differences in cognitive performance has important implications for developing a scientific basis for social and medical prevention, education and intervention strategies. Anatomy is unlikely to be readily affected by behavioral or pharmacologic treatment. By contrast, physiologic states such as measured by CBF, ALFF, ReHo and BOLD activation, can be changed within seconds, and it is easier to conceive of treatments that can affect resting-state CBF for sustainable durations. Our results suggest questions for future investigation. For example, current methods for rehabilitation of brain dysfunction emphasize activation of task-related brain systems. Our findings that lower resting CBF and increased resting state connectivity are associated with better performance suggest that emphasis should also be placed on training in deactivation of task-relevant regions in the absence of a task. Indeed, our results may offer an avenue for future scientific probing of the benefits of procedures such as meditation, which emphasize relaxation associated with the absence of goal-oriented behavior and results in reduced default-mode activity (Brewer et al. 2011; Hasenkamp and Barsalou 2012). That both greater volume and gray matter density of brain and lower basal metabolic rate are associated with cognitive abilities is consistent with preservation of tissue at low energy consumption as the “holy grail” for optimal organ function.

## ACKNOWLEDGMENTS

We thank the participants of the Philadelphia Neurodevelopmental Cohort and the members of the Recruitment, Assessment, Neuroimaging and Data Teams whose contributions made this project possible. This work was supported by NIH grant MH107235, MH089983, MH096891, MHP50MH06891, R01MH113550, R01MH112847, the Dowshen Neuroscience fund, and the Lifespan Brain Institute of Children’s Hospital of Philadelphia and Penn Medicine, University of Pennsylvania.

## AUTHOR CONTRIBUTIONS

R.E.G. and R.C.G. conceived the project, designed the study, guided data analysis, interpreted the results and wrote the manuscript. M.A.E., R.V. and J.A.D. designed the imaging protocol and participated in data analysis, R.C.G., E.R.B., T.M.M., A.F.G.R., A.P. and K.R. analyzed data, made figures and wrote sections of the manuscript. T.D.S., D.R.R., D.H.W., R.V., C.D. and E.D.G. guided image processing and interpretation of results. W.B.B. and R.T.S. guided statistical analysis. All authors reviewed and contributed to the write-up of the manuscript.

